# A staphylococcal cyclophilin carries a single domain and unfolds via the formation of an intermediate that preserves cyclosporin A binding activity

**DOI:** 10.1101/511048

**Authors:** Soham Seal, Soumitra Polley, Subrata Sau

## Abstract

Cyclophilin (Cyp), a peptidyl-prolyl *cis*-*trans* isomerase (PPIase), acts as a virulence factor in many bacteria including *Staphylococcus aureus*. The enzymatic activity of Cyp is inhibited by cyclosporin A (CsA), an immunosuppressive drug. To precisely determine the unfolding mechanism and the domain structure of Cyp, we have investigated a chimeric *S. aureus* Cyp (rCyp) using various probes. Our limited proteolysis and the consequent analysis of the proteolytic fragments indicate that rCyp is composed of one domain with a short flexible tail at the C-terminal end. We also show that the urea-induced unfolding of both rCyp and rCyp-CsA is completely reversible and proceeds via the synthesis of at least one stable intermediate. The secondary structure, tertiary structure, and the hydrophobic surface area of no intermediate are fully identical to those of other intermediate or the related native protein. Further analyses reveal no loss of CsA binding activity in rCyp intermediate. The thermodynamic stability of rCyp was also significantly increased in the presence of CsA, recommending that this protein could be employed to screen new CsA derivatives in future.

## Introduction

The cyclophilins (EC: 5.2.1.8) represent a family of highly conserved peptidyl-prolyl *cis/trans* isomerase (PPIase) enzymes those are expressed by most living organisms, and some giant viruses [1–5]. These proteins control protein folding by catalyzing the *trans* to *cis* isomerization of the peptidyl bonds those precede proline residues. These enzymes also influence numerous other cellular processes including protein trafficking, transcription, cell differentiation, apoptosis, protein secretion, T-cell activation, and signal transduction. In addition, these folding catalysts play critical roles in developing cardiovascular diseases, rheumatoid arthritis, viral infections, cancer, diabetes, sepsis, asthma, aging, neurodegenerative diseases, and microbial infections [2, 3, 6–11]. The catalytic activities of the cyclophilins are typically inhibited by cyclosporin A (CsA), a cyclic peptide harboring eleven amino acid residues [1]. A ternary complex, formed by the association of CsA-cyclophilin complex with calcineurin, prevents the dephosphorylation of the transcription factor NF-AT that, in turn, blocks the expression of cytokines from T-lymphocytes [12–14]. The reduction of T-cell activity by CsA has made it extremely useful in clinics, particularly for preventing the graft rejection after organ and bone marrow transplantation [2]. However, the severe side effects of CsA have restricted its use [15] and promoted to develop many CsA analogs with no immunosuppressive activity [2, 10, 11, 16–18]. Some of these CsA analogs though yielded promising results have not been approved yet.

Cyclophilins, located in the cytosol and membrane or cell organelles, are composed of either single domain or multiple domains [1–3, 19]. The catalytic domains of cyclophilins have a β-barrel conformation that is constituted with eight anti-parallel β-strands, two-three α-helices, and several connecting loops [1, 20, 21]. Their hydrophobic active sites are made using the side-chains of amino acid residues from most of the β-strands and loops. While thirteen residues are required for binding CsA [1, 20–22], eleven residues in the active site are involved in the binding of a tetrapeptide substrate [1, 23]. Of the residues, nine residues are used by both CsA and substrate for binding.

The linear polypeptide chains synthesized in the living cells become functional only when these molecules are folded into proper three-dimensional forms. To experimentally understand the mechanism of protein folding, native/denatured forms of proteins are gradually unfolded/refolded followed by monitoring their conformational changes using a suitable probe [24, 25]. The unfolding/refolding study usually indicates whether the folding of a protein occurs by a two-state or by a multi-state mechanism through the generation of no or multiple intermediates. Additionally, unfolding studies intimate about the stability of proteins in the presence of ligands or mutations [26–32]. Furthermore, such studies have greatly influenced many biotechnological fields including drug discovery [33–39]. The three-dimensional structures of the cyclophilins are conserved but their amino acid sequences are not identical [1–3], indicating that the unfolding mechanism of these proteins may be different. Thus far, unfolding mechanisms of only a few cyclophilins [40–43] were studied though these proteins were considered as the promising drug targets [2, 3, 10, 11].

*Staphylococcus aureus*, a pathogenic bacterium, harbors a putative cyclophilin (SaCyp)-encoding gene that is not induced by stress [44] or required for its growth [45]. The amino acid sequence of SaCyp shares adequate homology with those of cyclophilins from other organisms [42, 46]. A homology modeling study indicated the presence of a CsA binding site in SaCyp. Various experimental studies have collectively suggested that SaCyp is a PPIase, exits as a monomer, binds CsA, and plays a role in the *S. aureus*-mediated virulence [42, 46]. However, the catalytic activity of SaCyp later appears to be not critical for its pathogenesis [47]. The appearance and dissemination of the multi-drug resistant *S. aureus* strains across the globe today have necessitated the discovery of novel antistaphylococcal agents [48–50]. The structural and unfolding data of SaCyp may expedite the discovery of novel inhibitors (including new CsA derivatives) those would, in turn, be useful not only for treating staphylococcal infections but also other diseases. Recently, a study has indicated that the drug-bound or drug-unbound form of SaCyp unfolds via the generation of one intermediate in the presence of guanidine hydrochloride (GdnCl) [42]. In addition, there was a significant stabilization of SaCyp in the presence of CsA [42]. Proteins are sometimes denatured by a different pathway in the presence of different unfolding agent [51–54]. Thus far, the unfolding mechanism and stability of SaCyp have not been verified using other denaturants. The structural and functional properties of GdnCl-made SaCyp/SaCyp-CsA intermediate are also currently not known with certainty. Moreover, the predicted domain structure of SaCyp [42] has not been confirmed by any biochemical study. Herein, we have studied the domain structure and urea-induced unfolding of a recombinant *S. aureus* Cyp (rCyP) [42] using various probes. Our limited proteolysis data indicate that rCyp is a single-domain protein with a flexible tail at its C-terminal end. The urea-induced equilibrium unfolding of both rCyp and rCyp-CsA occurred via the synthesis of at least one stable intermediate. None of the intermediates has the properties of a molten globule [55]. Of the intermediates, rCyp intermediate has nearly full CsA binding activity.

## Materials and Methods

### Materials

Many materials including acrylamide, anti-his antibody, alkaline phosphatase-goat anti-mouse antibody, ANS (8-anilino-1-naphthalene sulfonate), bis-acrylamide, chymotrypsin, CsA, Phenylmethane sulfonyl fluoride (PMSF), isopropyl β D-1-thiogalactopyranoside (IPTG), proteinase K, protein marker, trypsin, and urea were used in the present study. We bought these materials from different companies such as Sigma, SRL, and Merck. rCyp was purified by a standard procedure as earlier reported [42].

### Basic protein techniques

The basic protein methods, namely, protein estimation, Western blotting, SDS-PAGE, and staining of polyacrylamide gel, were performed as reported [42, 56–58]. The theoretical mass of monomeric rCyp was determined by analyzing its sequence with ProtParam (web.expasy.org), a computational tool. The molar concentration of rCyp was estimated using both its theoretical mass and content in buffer B [42]. To produce rCyp-CsA, we incubated 10 μM rCyp in buffer B with 20 μM CsA for 30 min at 4°C [42].

### Functional investigation

The PPIase activity of rCyp (120 nM) was evaluated by RNase T1 (ribonuclease T1) refolding assay as reported [29, 42]. Previously, our modeling study indicated that one *S. aureus* Cyp molecule binds to one molecule of CsA [42]. Considering similar interaction between rCyp and CsA, the related equilibrium dissociation constant (*K*_d_) was estimated by a standard method [42] with minor modifications. Briefly, the intrinsic Trp fluorescence spectra (*λ*_ex_ = 295 nm and *λ*_em_ =300-400 nm) of rCyp (2 μM) in the presence of varying concentrations (0 - 4 μM) of CsA were recorded using a fluorescence spectrophotometer. The fluorescence intensity values (at *λ*_max_ =343 nm), extracted from the spectra, were rectified by deducting the related buffer fluorescence and by adjusting for volume changes. Lastly, the *K*_d_ value was estimated by fitting of the fluorescence data to a standard equation from GraphPad Prism (GraphPas Software Inc.).

### Limited proteolysis

To know whether rCyp carries any domain, limited proteolysis of this protein was separately executed by different proteolytic enzymes using standard methods [58, 59]. Briefly, a buffer B [42] solution carrying rCyp (10 μM) and an enzyme (0.025 - 0.1 μM) was incubated at ambient temperature. At different time points, an aliquot (50 µl) was pulled out and mixed with an SDS gel loading dye [56]. All of the aliquots were boiled prior to their resolution by a Tris-glycine SDS-13.5% PAGE. After staining with Coomassie brilliant blue, the photograph of acrylamide gel was captured as stated [58].

To determine the molecular masses of rCyp fragments, a MALDI-TOF analysis (Bruker Daltonics, Germany) was performed mostly as stated earlier [59]. Briefly, rCyp was exposed to a proteolytic enzyme for 10-20 min followed by the termination of reaction using PMSF at a final concentration of 0.5 mM. To inactivate the enzyme, the reaction mixture was incubated with benzamidine sepharose for 30 min. The supernatant collected after centrifugation was dialyzed against a 20 mM NH_4_HCO_3_ containing buffer for 4 h at 4°C. Finally, the supernatant obtained after centrifugation of the dialyzed sample was mixed with an equal volume of sinapinic acid. After drying the mixture on a sample plate, it was analyzed by an MALDI-TOF mass spectrometry. The yielded m/z spectra were used to calculate the molecular masses of the rCyp fragments using a standard method [60].

### Spectroscopic observation

To know about the different structural elements of rCyp and rCyp-CsA in buffer B [42], the ANS fluorescence (λ_ex_ / λ_em_ = 360 / 400-600 nm), intrinsic tryptophan (Trp) fluorescence (λ_ex_ / λ_em_ = 295 / 300-400 nm), near-UV circular dichroism (CD) (250-320 nm), and far-UV CD (200-260 nm) spectra of these proteins were recorded essentially as described before [42, 51, 58]. We used 25 μM protein for the near-UV CD spectroscopy and 10 μM protein for the far-UV CD or the fluorescence spectroscopy. The path length of cuvette in the near-UV CD spectroscopy was 5 mm, whereas that in the far-UV CD spectroscopy was 1 mm. The ANS concentration used in the study was 100 μM. The fluorescence or CD intensity values were rectified by subtracting the reading of buffer from the reading of buffer carrying protein.

### Unfolding and refolding of proteins

To study the unfolding pathway of rCyp and rCyp-CsA, these proteins (10 μM each) were exposed to varying concentrations (0-8 M) of urea for ~18 h at 4°C as stated [51, 58]. Protein aliquots were always treated with the freshly prepared urea solution. To understand the effects of denaturant on the different structures of proteins, the ANS fluorescence, intrinsic Trp fluorescence, and the CD spectra of the urea-treated/untreated proteins were recorded as described above. The spectroscopic signals were corrected by deducting the reading of urea containing buffer from the reading of the same buffer carrying protein.

To check whether the proteins denatured by 7-8 M urea can refold upon removal of urea, they were dialyzed against buffer B [42] prior to the recording of their Trp fluorescence spectra as described [51]. The spectra of equal extent of both native and unfolded proteins were also recorded for comparison. To see whether the refolded rCyp is functional, we performed RNase T1 refolding as stated [42].

### Transverse urea gradient gel electrophoresis

The unfolding of rCyp and rCyp-CsA were also monitored by a standard transverse urea gradient gel electrophoresis (TUGE) with minor modifications [29, 51, 61]. Briefly, a gel having a 10 - 7% acrylamide gradient and a 0 - 8 M urea gradient was made using a 10% acrylamide solution and a 7% acrylamide solution containing 8 M urea. At 0 and 8 M urea, the concentrations of acrylamide were 10% and 7%, respectively. After turning the solidified gel 90º, protein (60 µg) in an SDS-less loading buffer [56] was loaded on its generated well. Electrophoresis and staining of the gel were performed using standard procedures [57].

### Analysis of unfolding data

To gather clues about the unfolding pathways and the stabilities of rCyp and rCyp-CsA, the unfolding curves, produced using their spectroscopic and TUGE data, were fit to either the two-state (N ↔ U) or the three-state (N ↔I↔U) equation using GraphPad Prism as described [24, 25, 29, 58]. Of the yielded thermodynamic parameters, *C*_m_, urea concentration at the midpoint of unfolding transition (i.e. urea concentration when Δ*G* = 0), Δ*G*^W^, free energy change in the absence of urea, *m*, cooperativity parameter of unfolding, and ΔΔ*G*, the difference of free energy change between rCyp-CsA and rCyp, were considered in the study. The fraction of unfolded protein molecules was determined from the unfolding data using a standard method [29].

## Results

### Domain structure of rCyp

A modeling study previously indicated that the cyclophilin, encoded by *S. aureus*, could be a single domain protein [42]. To confirm this proposition, we have individually performed limited proteolysis [51, 58–60, 62] of rCyp with trypsin, chymotrypsin, and proteinase K. Each of these enzymes is computationally determined to have higher than ten cleavage sites, which are distributed along the entire sequence of rCyp (Fig. 1A). This protein will mostly remain insensitive to the above enzymes if it is really composed of only one domain. We have noted the generation of primarily one proteolytic fragment from rCyp at the initial stage of its cleavage with proteinase K (Fig. 1B). One major fragment was also made at the early period of digestion of rCyp with trypsin (Fig. 1C) or chymotrypsin (Fig. 1D). The proteinase K-, trypsin- and chymotrypsin-generated fragments are designated as fragment I, fragment II and fragment III, respectively. All of the fragments remained stable during the entire period of digestion. The intensities of the fragments were gradually increased with the increase of time of digestion. Their molecular masses are about ~2-3 kDa less than that of rCyp, indicating that the digestion occurred at one end or both ends of this protein in the presence of proteinase K. Conversely, fragment II was possibly originated due to the removal of any one end of rCyp. On the other hand, fragment III might have been generated due to the cleavage of the less chymotrypsin-sensitive peptide bonds formed by Leu, Met, and His residues (web.expasy.org/peptide_cutter) at the end(s) of rCyp. Our Western blot analyses show no interaction between the proteolytic fragments and anti-His antibody (Figs. 1E-1G), indicating the loss of polyhistidine tag from the N-terminal end of rCyp in the presence of the above enzymes.

**Fig. 1.**
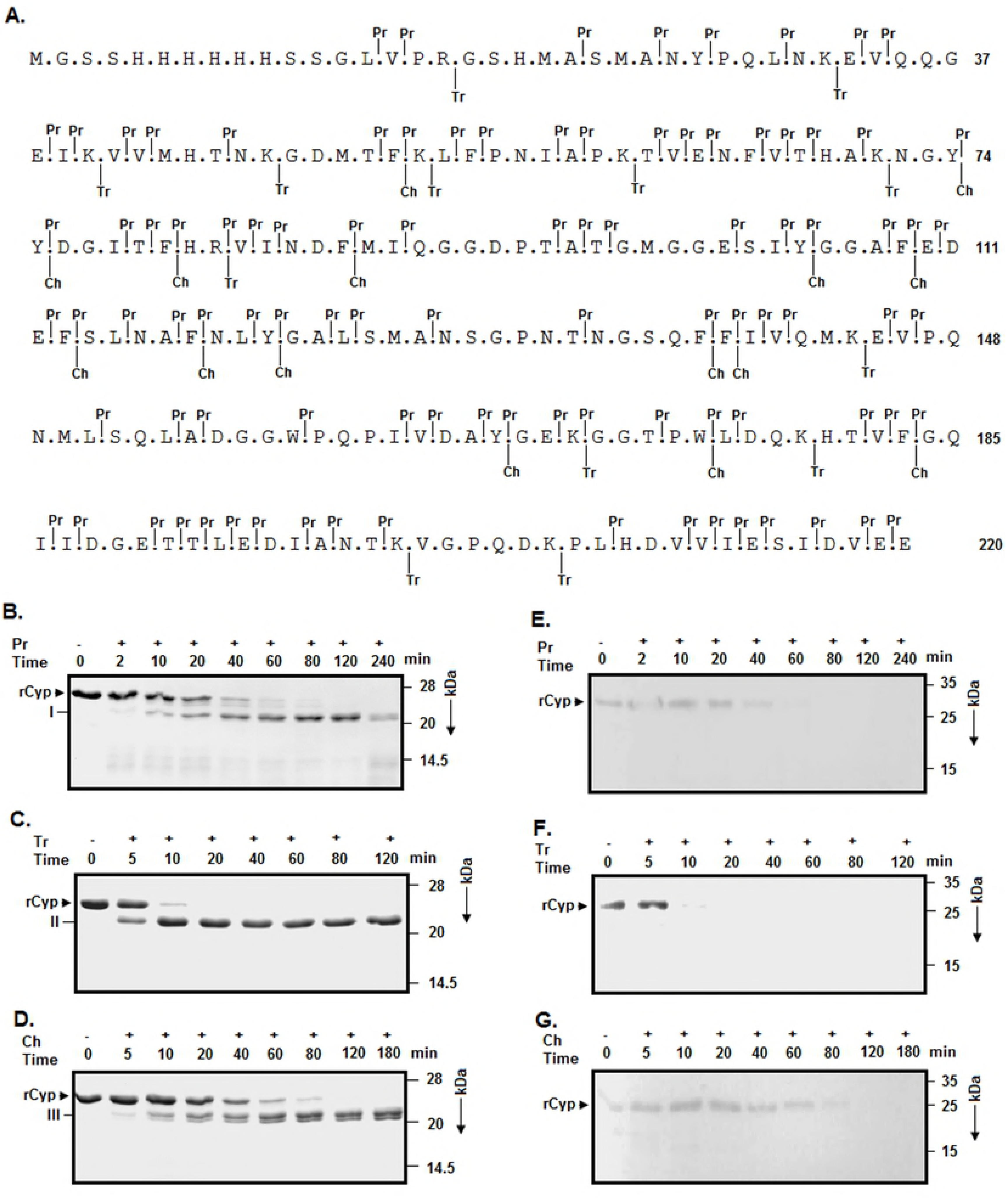
Characterization of rCyP by limited proteolysis. (A) The amino acid sequence of rCyp with the cut sites of chymotrypsin (Ch), trypsin (Tr), and proteinase K (Pr). The predicted cleavage sites, found out by a software tool [58], are shown at the top and bottom of the sequence using vertical lines. The polyhistidine tag of rCyp is composed of residues 1-23. The rCyp fragments, generated from the partial digestion of rCyp with the either of Pr (B), Tr (C), and Ch (D), were separated by different SDS-13.5% PAGE. Arrowhead and I-III represent the intact rCyp and the major rCyp fragments, respectively. (E, F, G) Western blotting experiment. The Pr-, Tr-, and Ch-generated rCyp fragments were analyzed using an anti-his antibody. We have mentioned the mass of each marker protein at the right-hand sides of the blot and gel images.

To find out the cut sites in rCyp, the masses of the above proteolytic fragments (I-III) were estimated using a MALDI-TOF mass spectrometry as described [59]. The m/z spectrum shows that there was a generation of two major peaks from rCyp digested with proteinase K (S1A Fig.). Conversely, trypsin (S1B Fig.)- or chymotrypsin (S1C Fig.)- digested rCyp resulted in largely one major peak as expected. As the two peaks obtained from the proteinase K-digested rCyp were fused with each other, the fragment I might be composed of two proteolytic fragments (designated as Ia and Ib) having a little difference in molecular mass. The single major peak originated from the trypsin-digested rCyp most possibly corresponds to fragment II. Similarly, the peak yielded from the chymotrypsin-cleaved rCyp might be due to fragment III. The molecular masses of the above rCyp fragments, calculated using the m/z spectral data (S1 Fig.), were found to vary from 21493.63 to 22050.73 Da (Table 1). Using the predicted cut site data of rCyp (Fig. 1A), different proteolytic fragments were generated followed by the determination of their masses using a computational tool (web.expasy.org/protparam). The rCyp fragments whose theoretical masses nearly matched with the experimental masses of fragments Ia, Ib, II, and III are composed of the amino acid residues Ser 23 to Val 218, Ser 23 to Glu 219, Gly 18 to Glu 220, and Ala 22 to Glu 220, respectively (Fig. 1A). Thus, five peptide bonds, made by the rCyp residues Arg 17 and Gly 18, Met 21 and Ala 22, Ala 22 and Ser 23, Val 218 and Glu 219, and Glu 219 and Glu 220, showed sensitivity to the proteolytic enzymes employed in the investigation. Of the susceptible peptide bonds, three bonds are in the polyhistidine tag carrying region of rCyp and the rest bonds are in the extreme C-terminal end of this enzyme (Fig. 1A). Collectively, both ends of rCyp might be exposed to its surface.

**Table 1.** Mass and composition of the proteolytic fragments.

| Enzyme | rCyp fragments | Mass of rCyp fragments<sup>A</sup> (Da) | Composition of rCyp fragments<sup>B</sup> |
| --- | --- | --- | --- |
| Proteinase K | Ia | 21493.63 | Ser23-Val218 |
| Proteinase K | Ib | 21564.43 | Ser23-Glu219 |
| Trypsin | II | 22050.73 | Gly18-Glu220 |
| Chymotrypsin | III | 21973.69 | Ala22-Glu220 |

<sup>A</sup>The masses of the rCyp fragments were estimated using MALDI-TOF data (S1 Fig.).
<sup>B</sup>The masses of fragments with the designated residue carrying regions, calculated by a
computational tool (web.expasy.org/protparam), are very close to those determined using MALDI-TOF data.

### Unfolding of proteins

The unfolding pathways of many proteins appeared dissimilar in the presence of different denaturants [49–51, 63]. Previously, both rCyp and rCyp-CsA in the presence of GdnCl were unfolded via the production of one intermediate [42]. To check whether the urea-induced unfolding of these proteins would follow the similar pathway, their far-UV CD, intrinsic Trp fluorescence, and ANS fluorescence spectra were separately recorded in the presence of 0 to 7/8 M urea (S2 Fig.) A monophasic curve is obtained for rCyp when the ellipticity values at 222 nm were plotted against the matching urea concentrations. Conversely, such a curve generated for rCyp-CsA was biphasic in nature (Fig. 2A). A monophasic curve for rCyp and a biphasic curve for rCyp-CsA were also obtained when we plotted their Trp fluorescence intensity (Fig. 2B) or the associated *λ*_max_ (Fig. 2C) values against the related urea concentrations. The *λ*_max_ values of both proteins were shifted to 350 nm when there was a saturation of fluorescence intensity. All of the biphasic curves show the transitions at ~1.5/2 - 2.75/3 M and ~5/5.5 – 7/7.5 M urea, respectively. Unlike the curves obtained using the CD and Trp fluorescence data, the curves, prepared using the ANS fluorescence intensity values of rCyp and rCyp-CsA, look very similar and possibly carry two transitions (Fig. 2D).

**Fig. 2.**
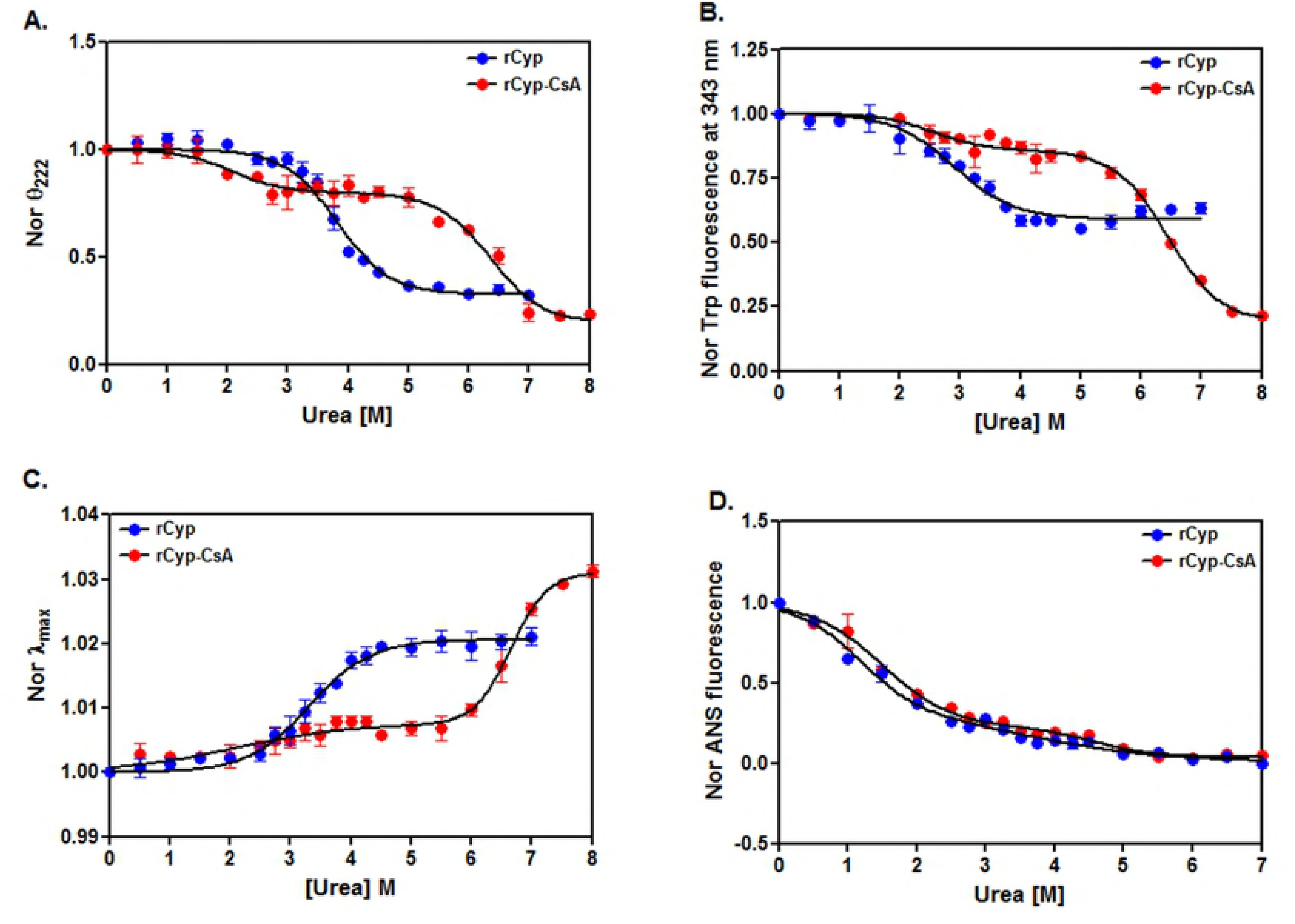
Unfolding studied by spectroscopic tools. (A) The ellipticity values of rCyp and rCyp-CsA at 222 nm, extracted from their far-UV CD spectra (S2 Fig.), were normalized and plotted against the corresponding urea concentrations as described [51]. (B) The intrinsic Trp fluorescence intensity values of rCyp (at 343 nm) and rCyp-CsA (at 341 nm), obtained from the respective spectra (S2 Fig.), were normalized and plotted as above. (C) The λ_max_ values of rCyp and rCyp-CsA, derived from the Trp fluorescence spectra (S2 Fig.), were similarly plotted. (D) The ANS fluorescence intensity values of rCyp, and rCyp-CsA at 480 nm, collected from the related spectra (S2 Fig.), were identically normalized and plotted. All lines through the spectroscopic signals denote the best-fit lines.

To verify the above unfolding data, we have also investigated the unfolding of rCyp and rCyp-CsA using transverse urea gradient gel electrophoresis [51], a biochemical probe. The migration of rCyp or rCyp-CsA across the urea gradient gel yielded an S-shaped protein band having nearly a clear transition region (Fig. 3). The rCyp-specific protein band shows a transition at ~3.25-4.25 M urea, whereas, that of rCyp-CsA results in a transition region at ~4.75-5.75 M urea, indicating that the initiation of the unfolding of drug-bound rCyp occurred at higher urea concentration. The faded transition region also suggests a slow unfolding reaction [59, 61].

**Fig. 3.**
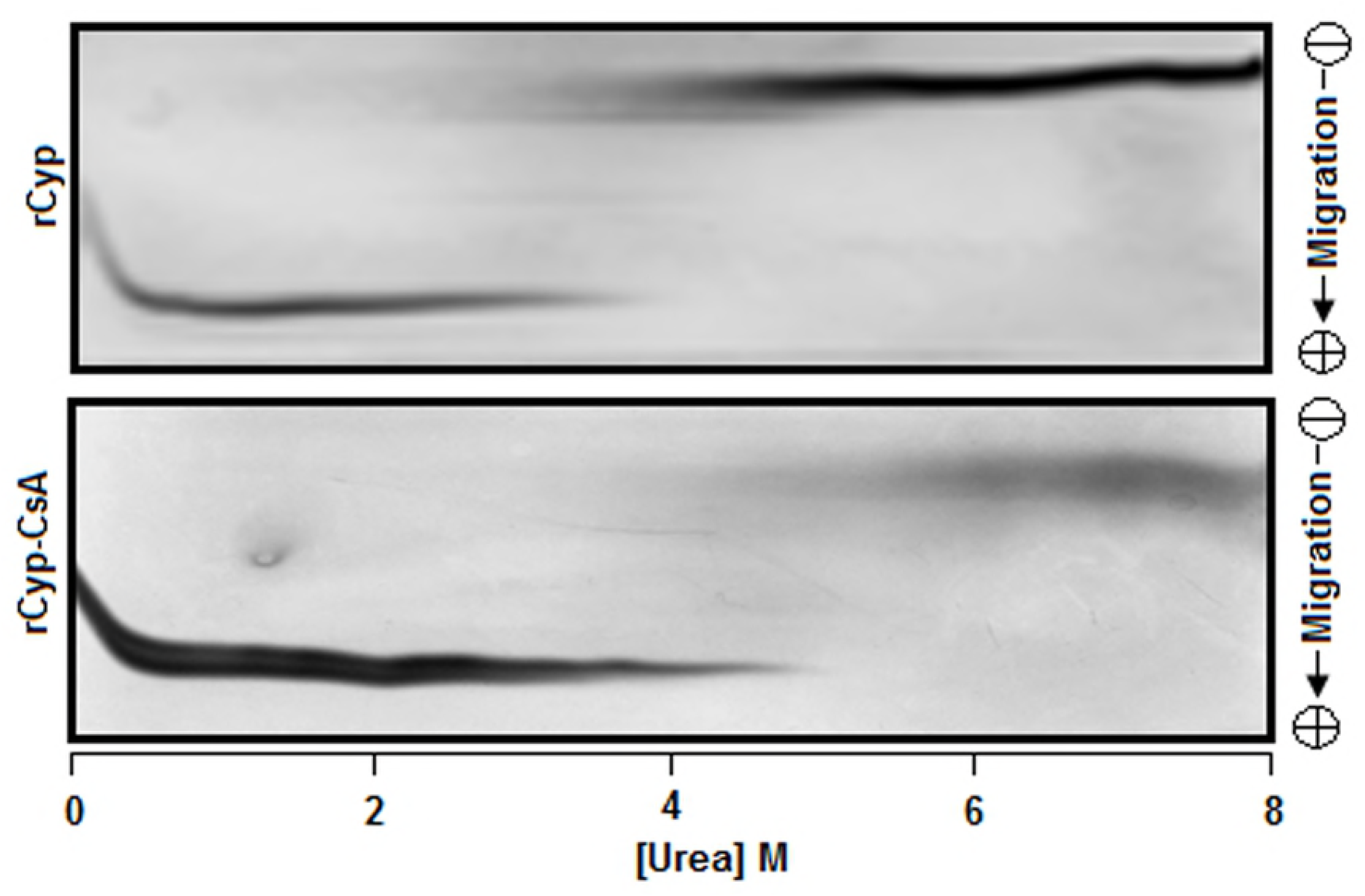
Transverse urea gradient polyacrylamide gel electrophoresis of proteins. Both rCyp and rCyp-CsA were separately analyzed by the Transverse urea gradient polyacrylamide gel electrophoresis as described [51, 59].

The reversibility of the unfolding reaction was checked by recording the Trp fluorescence spectra of the native, denatured, and the probable refolded forms of rCyp and rCyp-CsA as described [42]. We have observed that the Trp fluorescence spectra of the native protein and the related refolded protein have completely coincided with each other (Fig. 4A). Additional RNase T1 refolding assay reveals that there is nearly a complete restoration of the PPIase activity in the renatured rCyp (Fig. 4B). In sum, both rCyp and rCyp-CsA were unfolded by a reversible pathway in the presence of urea.

**Fig. 4.**
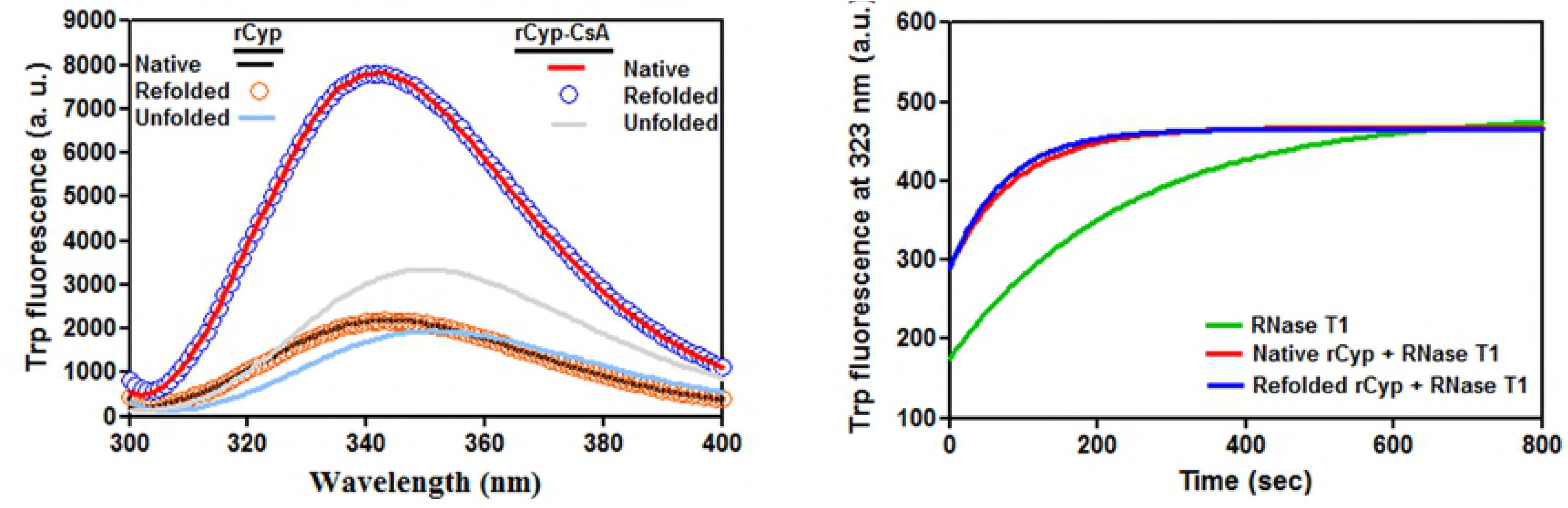
Refolding of the urea-exposed proteins. (A) The intrinsic Trp fluorescence spectra of the unfolded, native, and refolded rCyp or rCyp-CsA. (B) RNase T1 activity of refolded and native rCyp.

### Unfolding mechanism of the protein

To accurately determine the mechanism of the urea-induced unfolding of rCyp and rCyp-CsA, all of the unfolding curves were examined using different models [24, 25, 58]. Each rCyp-specific curve, generated using CD or Trp fluorescence signals, exhibited the best fitting with a two-state model [24]. The *C*_m_ values, obtained from the fitted CD (Fig. 2A), Trp fluorescence intensity (Fig. 2B) and the *λ*_max_ (Fig. 2C) data of rCyp, are 3.82±0.04 M, 2.91 ± 0.07 M, and 3.30±0.06 M urea, respectively. Conversely, the rCyp-specific curve, produced using ANS fluorescence signals, fit best to the three-state model [25] with the resulted *C*_m_ values of ~1.18 M and ~3.68 M urea (Table 2). Thus, the ANS fluorescence data suggest the formation of a rCyp intermediate at ~3 M urea (Fig. 5). Two additional pieces of evidence have supported the above proposal. The fractions of denatured rCyp molecules, estimated from both the CD (Fig. 2A) and Trp fluorescence data (Fig. 2B), were plotted against 0-7 M urea and the resulted curves did not coincide with each other (S3A Fig.). The non-overlapping of such curves indicates the formation of unfolding intermediate [25]. Secondly, the phase diagram [29, 64], used to know about the formation of the hidden unfolding intermediate of proteins, was developed by plotting the Trp fluorescence intensities of rCyp at 320 nm against its fluorescence intensities at 365 nm (S3B Fig.). The yielded non-linear plot again suggests the formation of rCyp intermediate(s) at 0-7 M urea.

**Fig. 5.**
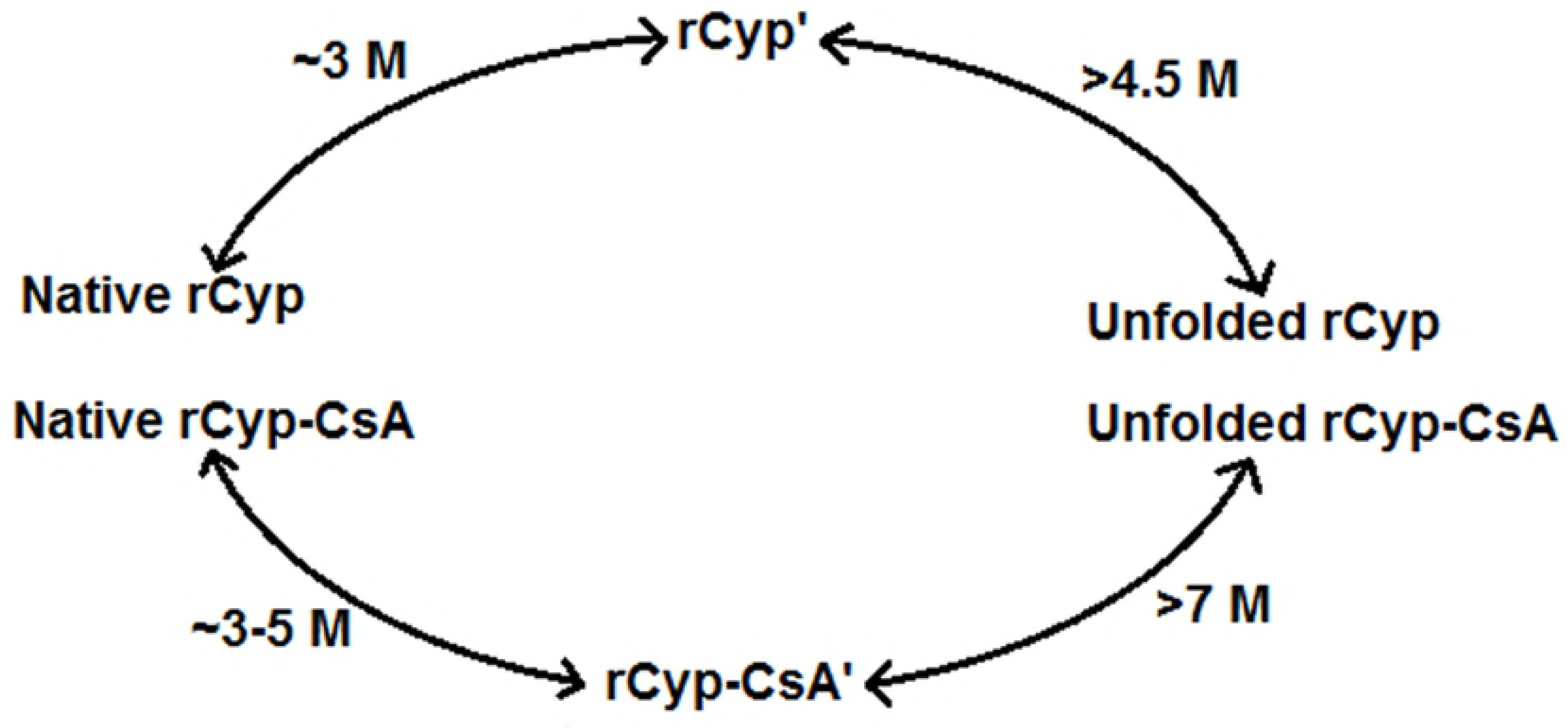
A graphic presentation of the urea-induced unfolding of rCyp and rCyp-CsA. The unfolding intermediates are denoted by rCyp’ and rCyp-CsA’.

**Table 2.** Thermodynamic parameters of protein unfolding.

| Protein | Assay method | Fitted equation | $C_m$ (M) | $m$ (kcal mol<sup>-1</sup>M<sup>-1</sup>) | $\Delta G^w$ (kcal M<sup>-1</sup>) | $\Delta\Delta G$ (kcal M<sup>-1</sup>) |
| --- | --- | --- | --- | --- | --- | --- |
| rCyp | ANS fluorescence | Three-state | 1.18±0.03<br>3.68±0.19 | 1.69±0.01<br>0.74±0.02 | 1.99±0.03<br>2.74±0.07 |  |
|  | TUGE | Two-state | 3.79±0.36 | 1.81±0.21 | 6.83±0.15 |  |
| rCyp-CsA | ANS fluorescence | Three-state | 1.51±0.02<br>4.61±0.21 | 1.33±0.03<br>1.70±0.19 | 2.01±0.12<br>7.82±0.53 | 0.55±0.10<br>1.12±0.08 |
|  | TUGE | Two-state | 5.53±0.56 | 1.44±0.19 | 7.92±0.27 | 2.81±0.02 |

The thermodynamic parameters were estimated from ANS fluorescence (Fig. 2D) and TUGE (Fig. 3) data using standard equations [24, 25].

All of the unfolding data of rCyp-CsA (Fig. 2), accumulated from our spectroscopic studies, fit best to a three-state model, clearly suggesting the formation of a rCyp-CsA intermediate in the presence of urea. While the *C*_m_ values yielded from the CD data of rCyp-CsA are 2.09±0.22 M and 6.37±0.09 M, those from its Trp fluorescence data are 2.58±0.17 and 6.5±0.05 M urea. Conversely, the *C*_m_ values estimated from the ANS fluorescence signals of rCyp-CsA are ~1.51 M and ~4.61 M urea, respectively. Jointly, a rCyp-CsA intermediate might have been generated at ~3-5 M urea (Fig. 5).

### Stability of the protein

A protein is usually stabilized when it binds a ligand [28, 29, 33, 36, 37, 39, 42]. To see whether the stability of rCyp is increased in the presence of CsA, the *C*_m_, *m*, Δ*G*^W^, and ΔΔ*G* values (Table 2), obtained from the ANS fluorescence data of rCyp and rCyp-CsA (Fig. 2D), were further analyzed as state above. The data show that the *C*_m_ values of rCyp-CsA are significantly higher than those of rCyp (all *p* values <0.05). The difference of free energy change between rCyp and rCyp-CsA (i.e. ΔΔ*G*) is more than ~0.5 kcal M^-1^ (Table 2). The thermodynamic parameters, determined by fitting the TUGE data (Fig. 3) with the two-state equation [24], are also presented in Table 2. The yielded Δ*G*^W^ and *C*_m_, values of rCyp-CsA were noted to be significantly higher than those of rCyp (all *p* values ≈ 0.03). The free energy change ΔΔ*G* between rCyp and rCyp-CsA is about 2.8 kcal M^-1^ (Table 2). Taken together, we suggest that the stability of rCyp is increased in the presence of CsA.

### Properties of unfolding intermediates

To confirm the generation of unfolding intermediates, the urea-exposed rCyp and rCyp-CsA were separately digested with trypsin as stated [58]. The yielded proteolytic patterns of proteins at ~0-1 M urea look different from those in the presence ~2-6 M urea (Fig. 6A). While new proteolytic fragments from rCyp appeared at ~3-6 M urea, those from rCyp-CsA are generated at ~2-6 M urea. The emergence of the additional proteolytic fragments might be due to the change of protein structure at the above urea concentrations. Thus, the data prove the production of unfolding intermediates from both proteins at moderately higher urea concentrations.

**Fig. 6.**
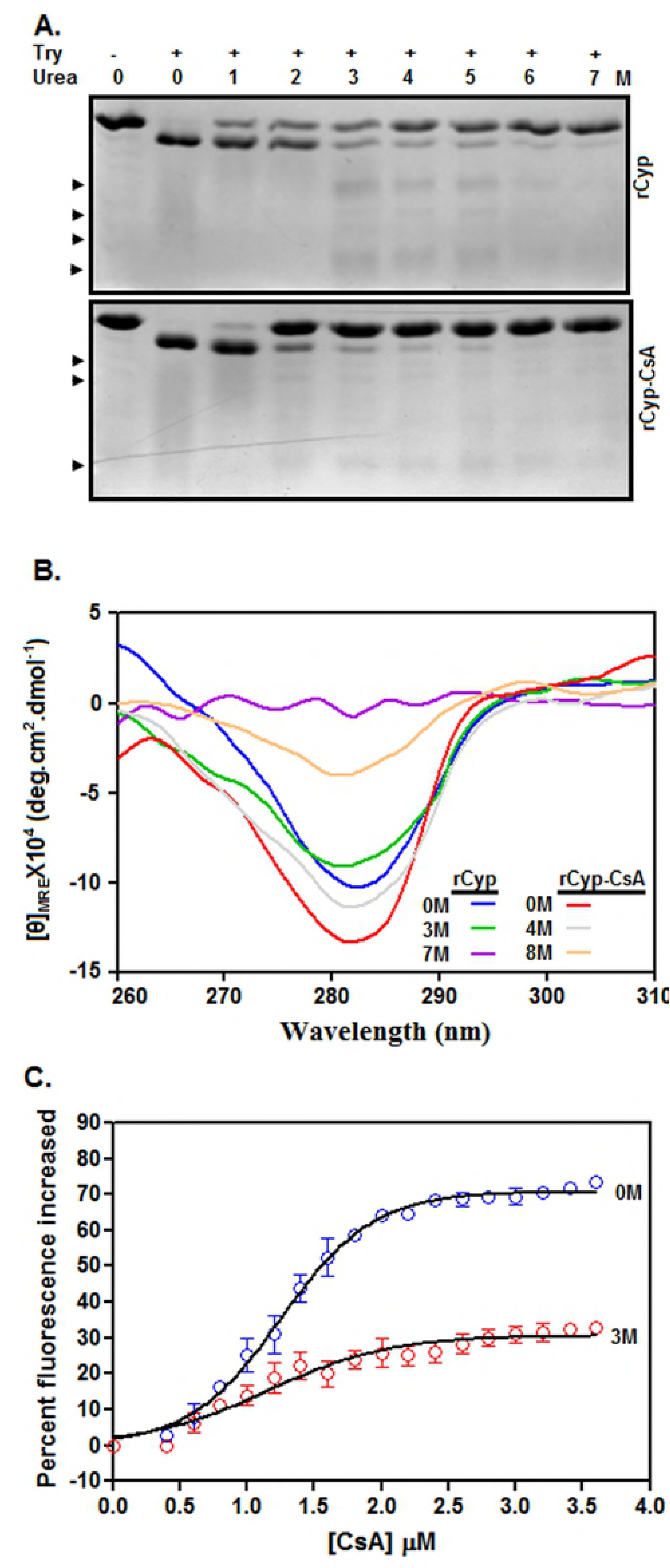
Properties of urea-made intermediates. (A) Analyses of the trypsin-generated fragments from rCyp and rCyp-CsA. Proteins were exposed to 0-7 M urea followed by their digestion with (+)/without (-) trypsin for 10 min at 25°C. All of the proteolytic fragments are resolved by SDS-13.5% PAGE. Arrowheads denote new protein fragments. (B) The near-UV CD spectra of rCyp and rCyp-CsA at the denoted concentrations of urea. The spectra were recorded using same concentrations of proteins. (C) Drug binding assay. The curves show the change of Trp fluorescence intensity of 0 M and 3 M urea-exposed rCyp (2 µM) in the presence of 0-3.5 µM Cyclosporin A (CsA).

The ellipticity value of rCyp at 222 nm was reduced about 4% when we enhanced the urea concentration from 0 M to 3 M urea (S4A Fig.), whereas, that of rCyp-CsA was dropped about 17% upon increasing the urea concentration from 0 M to 4 M urea. Therefore, both rCyp and rCyp-CsA intermediates are composed of sufficient extents of secondary structures.

The Trp fluorescence intensity of rCyp was decreased about 40% upon augmenting the urea concentrations from 0 M to 3 M (S4B Fig.). Conversely, there was a ~13% reduction of the Trp fluorescence intensity of rCyp-CsA when the urea concentration was enhanced from 0 M to 4 M. At the intermediate forming urea concentrations, the spectra of rCyp and rCyp-CsA are associated with the 4 nm and 2 nm red-shifted emission maxima, respectively. In sum, the tertiary structures of rCyp and rCyp-CsA intermediates are different from those of the native forms of these proteins.

The ANS fluorescence intensities of rCyp at 3 M and rCyp-CsA at 4 M urea, unlike their far-UV CD and Trp fluorescence intensities, are more than 70% less in comparison with those of the related proteins at 0 M urea (S4C Fig.). Therefore, the extent of the hydrophobic surface area in either intermediate is significantly less than that in the related native protein.

The near-UV CD values of rCyp at ~280-285 nm were slightly decreased when urea concentrations were augmented from 0 M to 3 M urea (Fig. 6B). On the contrary, the near-UV CD values of rCyp-CsA at ~280-285 nm were marginally reduced upon raising the urea concentrations from 0 M to 4 M. Collectively, both intermediates are not molten globules [55] as they retained sufficient extent of tertiary structures.

To verify whether rCyp intermediate is biologically active, we estimated the Trp fluorescence change of both 0 M and 3 M urea-equilibrated rCyp in the presence of different concentrations of CsA (Fig. 6C). The yielded *K*_d_ values for the rCyp and CsA interaction at 0 M and 3 M urea are 1.25 ±0.02 μM and 1.16±0.07 μM, respectively. Further analysis reveals no significant change of *K*_d_ value (*p*=0.16) upon changing the urea concentrations from 0 M to 3 M, indicating that rCyp intermediate did not lose any drug binding activity.

## Discussion

The present study has provided some seminal clues about the folding-unfolding mechanism and the domain structure of rCyp, a chimeric SaCyp harboring 220 amino acid residues [42]. Our limited proteolysis (Fig. 1) and the subsequent analyses (Table 1) have revealed that two rCyp ends carrying residues 1 to 22 and 218 to 220 are only susceptible to three proteolytic enzymes employed in the study. The rCyp region having residues 23 to 218 carries most of the cleavage sites of these enzymes (Fig. 1A). The absence of digestion in the internal rCyp region indicates the formation of a domain by the residues 23 to 218. Thus, the single domain structure of SaCyp proposed before on the basis of a modeling study data [42] was confirmed by our proteolysis results. However, such single domain structure is not unprecedented as the cyclophilins those have masses nearly similar to that of SaCyp are also shown to carry single domain capable of binding both the substrate and inhibitor [2, 3, 11, 19].

The proteolysis of Val 218 - Glu 219, and Glu 219 - Glu 220 peptide bonds (Fig. 1) indicates that the rCyp residues Val 218, Glu 219, and Glu 220, corresponding to SaCyp residues Val 195, Glu 196, and Glu 197, might be exposed to its surface. Our computational examination of the model SaCyp structure [42] reveals that four C-terminal end residues, Asp 194, Val 195, Glu 196 and Glu 197, are not involved in the formation of any secondary structure and more than 20% exposed to its surface (data not shown). We have noted that the extreme C-terminal end of some SaCyp homologs, formed by three to four residues, are also not structured but adequately surface-exposed. The above data not only support our proteolysis data but also indicate that a short flexible region is attached to the C-terminal end of SaCyp domain. Thus far, no other single domain cyclophilin was reported to carry a short tail at the C-terminal end.

Our spectroscopic data have indicated that unfolding of rCyp or rCyp-CsA at 0-7/8 M urea proceeds via the synthesis of one stable intermediate (Fig. 5). The unfolding pathway of either protein in the presence of urea was fully reversible though there was the production of an intermediate (Fig. 4). The surface hydrophobicity, secondary structure, and the tertiary structure rCyp intermediate are not fully identical to those of rCyp-CsA intermediate (Fig. 6 and S4 Fig.). The structural properties of the intermediates also do not match with those of native proteins. Of the intermediates, the rCyp intermediate is formed at comparatively less concentration of urea (Fig. 5). Collectively, the number and type of non-covalent interactions responsible for stabilization of a protein structure [68, 69] are possibly not identical in the two intermediates.

The unfolding mechanism of rCyp and rCyp-CsA in the presence of urea (Fig. 5) matches with that of these proteins in the presence of GdnCl [42]. However, the GdnCl concentration needed to make the rCyp intermediate was higher than that required to synthesize rCyp-CsA intermediate. The structural properties of the urea-made intermediates are also not completely identical to those of the GdnCl-generated intermediates (Fig. 6). The GdnCl-made rCyp intermediate, unlike the urea-produced rCyp intermediate (Fig. 6B), has a little extent of tertiary structure (S5A Fig.). On the other hand, the GdnCl-made rCyp-CsA intermediate, like the urea-created rCyp-CsA intermediate, has retained some tertiary structure (S5A Fig.) but lost most of the hydrophobic surface area (S5B Fig.). In sum, neither the GdnCl-generated nor the urea-made intermediates have the properties of a molten globule [55].

The unfolding mechanism of drug-bound/unbound rCyp shows some similarity and dissimilarity with those of other drug-bound/unbound cyclophilins [40, 41, 43]. A yeast-encoded cyclophilin (CPR3) in the presence of urea was unfolded by means of the creation of two structurally different intermediates [43]. Of the CPR3 intermediates, the intermediate formed at relatively less urea concentration has the characteristics of a molten globule [55]. Like rCyp, LdCyp, a cyclophilin synthesized by *Leishmania donovani* [41], was unfolded by a three-state mechanism in the presence of GdnCl. However, the LdCyp intermediate, unlike rCyp intermediate, has the properties of a molten globule [55]. On the other hand, the urea-induced unfolding of a mycobacterial cyclophilin (PpiA) or its drug-bound form occurred via the formation of an intermediate [40]. Currently, little is known about the biological activities of CPR3, LdCyp, and PpiA intermediates. Conversely, our studies for the first time have indicated that the drug binding activities of the urea-made rCyp intermediate and native rCyp are nearly similar (Fig. 6C). The GdnCl-made rCyp intermediate also retained about 25% of the total drug binding activity of native rCyp (S5 Fig.).

The complete retention of drug binding activity in the rCyp intermediate (Fig. 6C) implies no significant alteration of the three-dimensional structure of the cyclosporin A binding site in the presence of 3 M urea. Our previous studies showed that the putative cyclosporin A binding site in SaCyp is primarily located in the regions harboring residues ~Arg 59 to Phe 116 and ~Trp 152 to His 157 [42]. Therefore, the structural change noted in rCyp intermediate (Fig. 2) possibly have occurred at regions carrying residues ~Ala 2 to His 58, ~Ile 117 to Pro 151, and ~Thr 158 to Glu 197. Besides Trp 152, SaCyp carries another Trp residue at position 136 [42]. The former Trp residue is relatively more exposed on the surface of SaCyp (data not shown). As Trp 152 is conserved and indispensable for binding CsA [1], there might be little change of the structure around this residue in the rCyp intermediate. The altered Trp fluorescence intensity and emission maxima of the rCyp intermediate (Figs. 2B and 2C), therefore, suggests a structural change around Trp 136. Additional studies are needed to prove the urea-induced structural alteration around Trp 136 with certainty.

Many promising CsA analogs with no immunosuppressive activity were discovered and suggested to be useful in treating various diseases [2, 16–18, 70]. As these analogs did not yield encouraging results in the clinical trials [10, 11], screening or synthesis of additional CsA analogs should be continued on a priority basis. An inhibitor can be easily screened against a drug target if the binding of the former increases the midpoint of unfolding transition (or the stability) of the latter [33–39]. Several chemical denaturation-based assay systems were reported to screen the drug molecules against various drug targets including PPIase enzymes [28, 34, 35, 37, 39]. Our present (Table 2) and previous [42] unfolding results demonstrated the significant increment of the stability of rCyp in the presence of CsA. The urea-induced unfolding of a mycobacterial cyclophilin also reported its stabilization by CsA [40]. Collectively, an unfolding-based assay system could be developed using SaCyp or rCyp for screening new CsA analogs in the future.

## Conclusions

Our investigations have provided invaluable clues about the basic structure and the folding-unfolding mechanism of SaCyp, an *S. aureus*-encoded cyclophilin involved in pathogenesis. We noted that rCyp, a recombinant SaCyp, is a single-domain protein with a short tail at its C-terminal end. Additionally, rCyp unfolds via the formation of an intermediate in the presence of urea. The rCyp intermediate lacks the characteristics of a molten globule and also shows little loss of CsA binding activity. The unfolding of the CsA-bound rCyp also similarly occurred in the presence of urea. The stability data of rCyp seems to be applicable in the discovery of new CsA derivatives in the future.

## Acknowledgments

We thank Mr. S. Biswas, and Mr. M. Das for their excellent technical support.

